# Functional genomic signatures predict microbial culturability across the tree of life

**DOI:** 10.1101/2025.08.18.670795

**Authors:** Iyanu Oduwole, Ashley Babjac, Taylor M. Royalty, Mattie Hibbs, Karen G. Lloyd, Scott Emrich, Andrew D. Steen

## Abstract

Most microbial taxa on Earth remain uncultivated, limiting our ability to study their physiology, ecology, and roles in environmental processes. Although metagenome-assembled genomes (MAGs) have expanded access to uncultured phylogenetic diversity, the functional basis for culturability remains poorly understood. Here, we analyze the 52,515 MAGs from the Genomes from Earth’s Microbiomes (GEM) catalog to test two hypotheses: 1) genomes from uncultured microbes encode more functionally novel genes than those from cultured taxa, and 2) specific genomic features are systematically associated with culturability across phyla. To assess functional novelty, we aligned predicted proteins to SwissProt and measured sequence dissimilarity to the nearest curated homolog. We find that uncultured MAGs, particularly among Archaea, harbor substantially more divergent proteins. To identify genomic traits predictive of culturability, we combined pathway-level enrichment with LASSO regression and permutation-based feature importance. Cultured MAGs were consistently enriched in Clusters of Orthologous Groups (COG) pathways related to vitamin and cofactor biosynthesis (e.g., thiamine, folate, B12), energy metabolism (e.g., TCA cycle), and CRISPR-Cas systems—functions often depleted in uncultured counterparts. LASSO models identified a subset of these pathways as strong predictors of cultured status even in poorly sampled phyla, suggesting conserved genomic signatures of culturability. In contrast, pathways such as purine biosynthesis and NADH dehydrogenase were associated with uncultured lineages, highlighting potential barriers to cultivation. These results 1) demonstrate the great functional novelty of uncultured microbes, potentially offering unprecedented opportunities for discoveries of novel function, and 2) identify metabolic traits associated with culturability to inform future cultivation strategies.

**Importance:** The vast majority of microbes are uncultured, which means they have never been characterized under laboratory conditions. We showed that genomic sequences of uncultured microbes have less similarity to characterized proteins compared to cultured microbes, revealing that there may be fundamental biological reasons why they are not cultured. We also showed that certain metabolic pathways, such as those related to vitamin and cofactor biosynthesis, can predict the ability of microbes to grow under laboratory conditions, and these pathways are abundant in highly cultured phyla, indicating how metabolic pathways can influence cultivation strategies.

## Introduction

Most microbial cells on Earth belong to taxa that have never been grown in pure culture, despite decades of cultivation efforts (1–3), for reasons that are only partially understood (4–6). This cultivation gap limits our experimental access to microbial diversity and constrains efforts to characterize microbial physiology, ecological roles, and biotechnological potential.

One contributing factor may be uneven culturing efforts: some lineages may be underrepresented simply due to limited attempts to grow them from certain environments (1). A second explanation is that many microbes have complex or context-dependent growth requirements, including dependence on specific environmental cues or close associations with other taxa, which may not be easily replicated under standard lab conditions (7–11). Third, inherently slow-growing taxa may be outcompeted in enrichment cultures or may fail to reach detectable levels within typical incubation times (5, 12).

Although cultivation-independent techniques, particularly metagenome-assembled genomes (MAGs), have transformed our view of microbial diversity and expanded access to uncultured lineages (13, 14), the genomic basis of microbial culturability remains poorly understood. We reasoned that comparing cultured and uncultured genomes could identify functional trends associated with culturability and highlight protein-coding novelty in lineages that resist cultivation. Using over 50,000 MAGs from the Genomes from Earth’s Microbiomes catalog (15), we tested the following two hypotheses.

First, we hypothesized that uncultured microbes encode more functionally novel genes than cultured taxa. To assess novelty, we aligned predicted protein sequences from GEM to protein sequences in SwissProt, a manually curated database of experimentally characterized proteins and measured amino acid sequence dissimilarity to the closest known homolog. Given that sequence divergence often reflects functional divergence (16, 17), we used this metric to infer protein-coding novelty relative to previously-characterized proteins. Second, we hypothesized that microbial culturability can be predicted from genomic content. Specifically, we examined the distribution of Clusters of Orthologous Groups (COG) pathways across genomes and used LASSO regression (18), a regularized modeling approach that selects a sparse set of predictive features, coupled with permutation-based feature importance (19), to identify genomic pathways most strongly associated with cultured status for each phylum.

Previous studies have shown that uncultured microbes are phylogenetically distinct from cultured lineages (1, 20). Our analysis extends these findings by showing that uncultured lineages —especially among Archaea— harbor extensive functional novelty, while cultured MAGs are consistently enriched in core metabolic pathways, particularly those involved in vitamins biosynthesis and energy metabolism. These patterns suggest that conserved genomic features may underlie culturability and may provide targets for improved cultivation strategies across phylogenetically diverse microbes.

## Results

### Cultured MAGs resemble cultured isolates

We analyzed 52,550 metagenome-assembled genomes (MAGs) from the GEM catalog. Of these, 14,620 were classified as “cultured” based on being classified at the species level with cultured isolates and sequence similarity to cultured isolates (>95% ANI, see Methods); the remaining genomes were classified as “uncultured.”

Cultured MAGs and cultured isolates exhibited similar levels of predicted *functional novelty*, defined as amino acid sequence dissimilarity to SwissProt proteins. Prior work suggests that protein sequences with ≥70% identity often retain similar biological functions (21). The comparable sequence similarity levels between cultured MAGs and isolates indicate that the process of sequencing DNA from the environment, assembly, and binning do not artificially inflate the apparent novelty (Fig. 1A-D).

**Fig 1:**
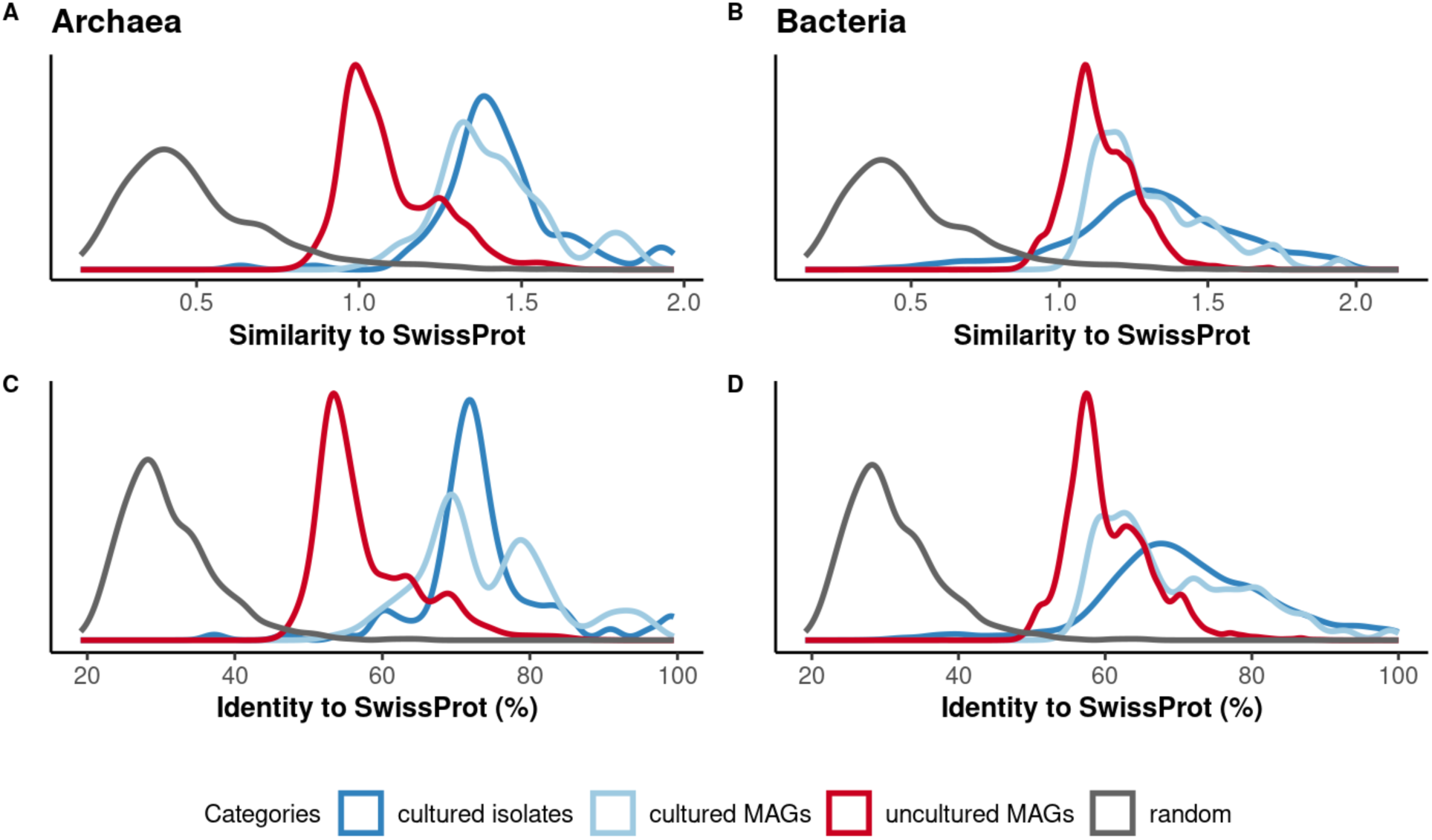
Similarity of predicted protein sequences in MAGs and isolates to SwissProt entries. Panels A and C show normalized bitscores and percent identities, respectively, for **archaeal** genomes; panels B and D show the same metrics for **bacterial** genomes. The distributions represent the average similarity of all predicted protein-coding genes to their closest match in the SwissProt. Random sequences (gray) represent simulated protein sequences with no known homology and serve as a negative control.

### Uncultured MAGs harbor substantially greater functional novelty

In contrast to the cultured MAGs, uncultured MAGs contained significantly more novel protein-coding genes than either cultured MAGs or isolates, based on both sequence identity and bitscore normalized to alignment length (Fig. 1A-D). To assess whether this increased novelty could result from spurious alignments, we compared all results to a control set of randomly generated sequences. All biological sequences—cultured and uncultured—awere substantially less divergent from SwissProt sequences than the random set (Fig. 1A-D), supporting the conclusion that the observed novelty reflects genuine biological differences rather than alignment noise.

Only 7% (2,725 out of 37,847) of predicted proteins in uncultured MAGs have exhibited more than 70% average amino acid identity with SwissProt homologs, compared to 42% (6,100 out of 14,620) in cultured MAGs and 51% (3,596 out of 7,035) in cultured isolates. These findings indicate that uncultured microbes encode extensive, unexplored protein diversity.

Uncultured archaeal MAGs were especially novel, exhibiting greater divergence from SwissProt proteins than uncultured bacterial MAGs relative to their respective cultured counterparts (Fig 1 A-D). This highlights uncultured archaeal lineages as particularly rich sources of uncharacterized functional diversity, although uncultured bacteria also have substantial functional novelty.

### Alignment metrics reflect genome source and quality

To evaluate whether genome quality contributes to these patterns, we examined alignment length distributions. Cultured MAGs had shorter alignment lengths than isolates (Fig S1a), likely reflecting lower sequencing depth or more fragmented assemblies (Fig S1a). As expected, uncultured MAGs had the shortest alignments, consistent with lower estimated completeness.

Across all genomes, alignment length was positively correlated with sequence identity— supporting the idea that more complete proteins yield more confident homology matches. By contrast, alignment length and identity were negatively correlated in random sequences (Fig S1b). Correlation strength was highest in isolates (R = 0.42), intermediate in cultured MAGs (R = 0.18) and lowest in uncultured MAGs (R = 0.14) (Fig. S1b). These findings further support the biological validity of our results.

Roughly 9% of predicted proteins across all MAGs (combining Bacteria and Archaea) had no detectable match to SwissProt proteins, even with a permissive *e*-value threshold (*e* <100). These unmatched genes may represent pseudogenes, misannotated open reading frames (ORFs), or real proteins that have no homologues in any database.

### Phylogenetic resolution predicts functional similarity

Bacterial MAGs were more frequently assigned to specific taxonomic ranks than archaeal MAGs, consistent with a higher proportion of cultured representatives among bacterial lineages (Fig. S2). In both domains, genomes assigned to more specific ranks (e.g., species) encoded proteins that were more similar to SwissProt entries (Fig. 2). This trend suggests that functional annotation is tightly linked to phylogenetic proximity to cultured taxa. A complementary analysis using the Agnostos HMM database yielded consistent results: the proportion of unclassified genes increased with taxonomic distance from cultured lineages (Fig S3). Genomes classified only at high taxonomic ranks (e.g., phylum) had the highest fraction of unannotated genes, whereas those resolved to the species level had the lowest.

**Fig 2:**
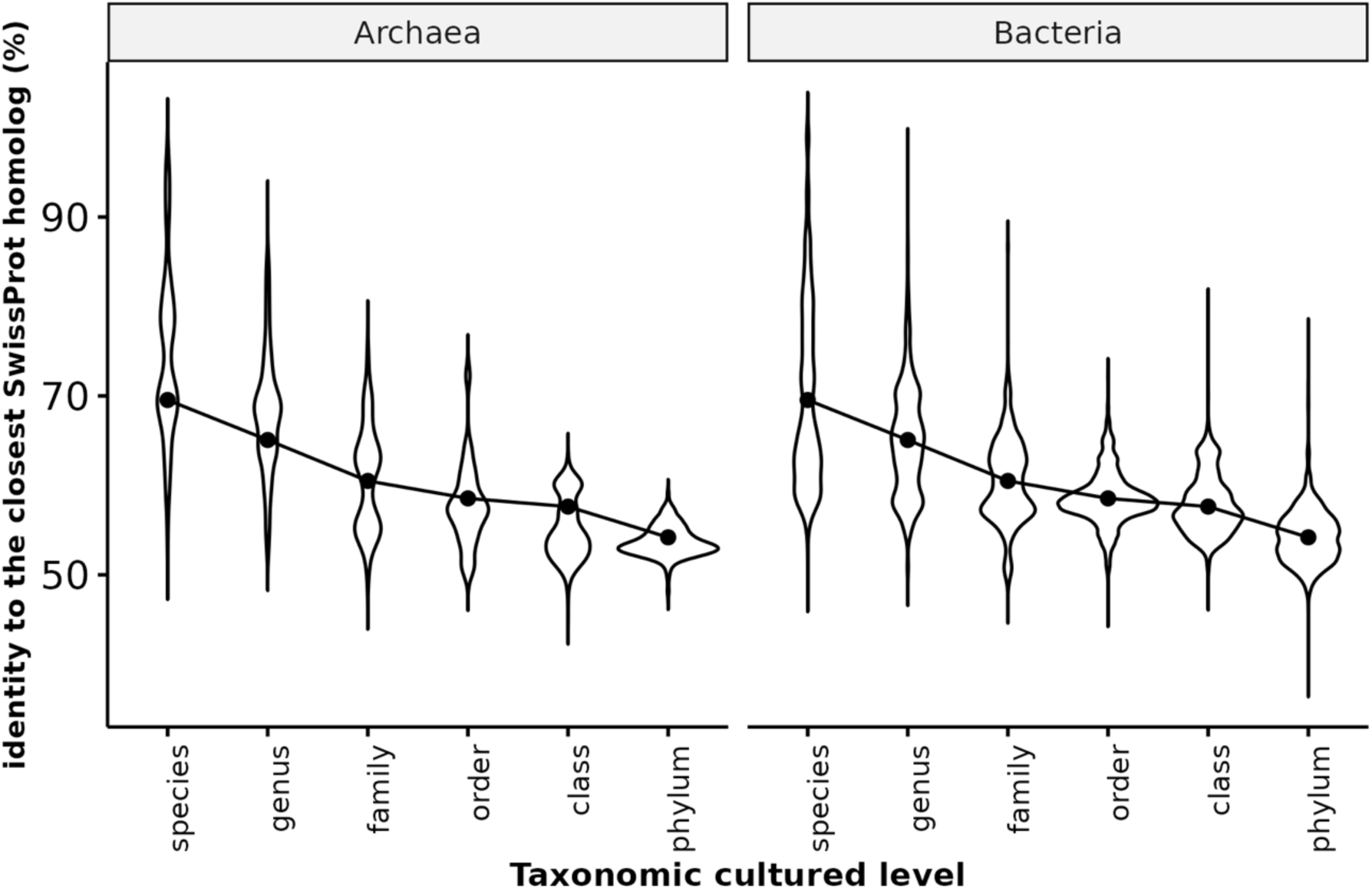
Similarity of predicted proteins in Archaea (left) and Bacteria (right) to their closest SwissProt homolog, stratified by taxonomic proximity to cultured isolates. For each metagenome-assembled genome (MAG) in the GEM catalog, the lowest shared taxonomic rank with a cultured isolate was determined. Protein similarity—measured as percent identity to the nearest SwissProt match—is shown as a function of this taxonomic proximity. Taxonomic rank assignments for GEM MAGs are based on the Genome Taxonomy Database (GTDB), as implemented by Nayfach et al. Cultured lineages were identified based on RefSeq annotations in the same GTDB release, excluding genomes derived from single-cell amplification or other metagenome-assembled genomes (MAGs or SAGs).

### Cultured microbes are enriched in certain metabolic pathways

To identify potential genomic features associated with culturability, we compared COG pathway enrichment in cultured versus uncultured MAGs, initially focusing on comparisons within individual phyla. We restricted this analysis to 28 phyla with sufficient numbers of both cultured and uncultured representatives, each containing at least one COG pathway significantly enriched in cultured or uncultured MAGs as determined by Fisher’s exact test.

Every COG pathway was significantly enriched in cultured MAGs in at least one phylum, and 46 out of 60 COG pathways were also enriched in uncultured MAGs in at least one phylum (Fig 3A). In most phyla, cultured MAGs were enriched for pathways involved in cofactor and vitamin biosynthesis, energy metabolism, and the CRISPR-Cas adaptive immunity. By comparison, uncultured MAGs were enriched for pathways such as NADH dehydrogenase, iron-sulfur biosynthesis, and asparagine biosynthesis (Fig 3A). These differences suggest that uncultured microbes may rely on alternative metabolic strategies, reduced biosynthetic capabilities, or niche-specific adaptations that hinder growth under laboratory conditions.

**Fig 3a:**
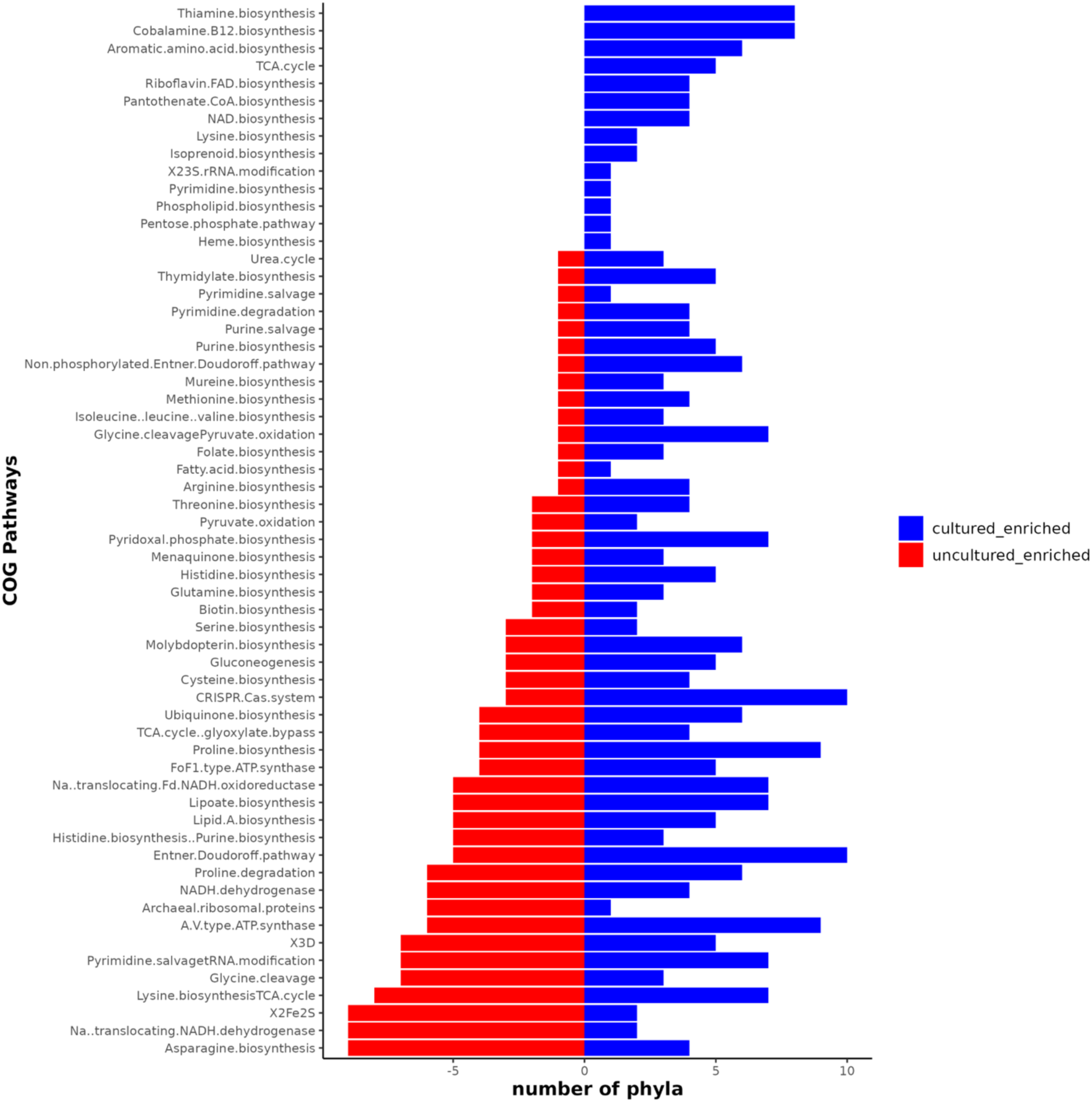
COG pathways that are enriched in cultured and uncultured MAGs (P < 0.005, fisher’s exact test). Pathways related to co-factors, vitamins, energy production, and CRISPR Cas are over-represented in cultured MAGs.

We further identified 15 pathways that were enriched exclusively in cultured MAGs and in no uncultured MAGs across all phyla (Fig. 3B). These pathways may serve as genomic markers of culturability. Many were involved in cofactor biosynthesis—for example, thiamine biosynthesis was enriched in cultured members of 7–8 phyla including *Bacteroidota*, *Actinobacteriota*, and *Proteobacteria*. Cobalamin (B12) biosynthesis was enriched in *Firmicutes*, *Verrucomicrobiota*, *Cyanobacteriota*, and *Proteobacteria*. The TCA cycle was enriched in cultured members of the *Proteobacteria*, *Desulfobacterota*, *Fusobacteriota*, and *Firmicutes* groups. These findings suggest that a genome’s ability to support core metabolic processes and cofactor biosynthesis may promote microbial growth and survival under axenic laboratory conditions.

**Fig 3b:**
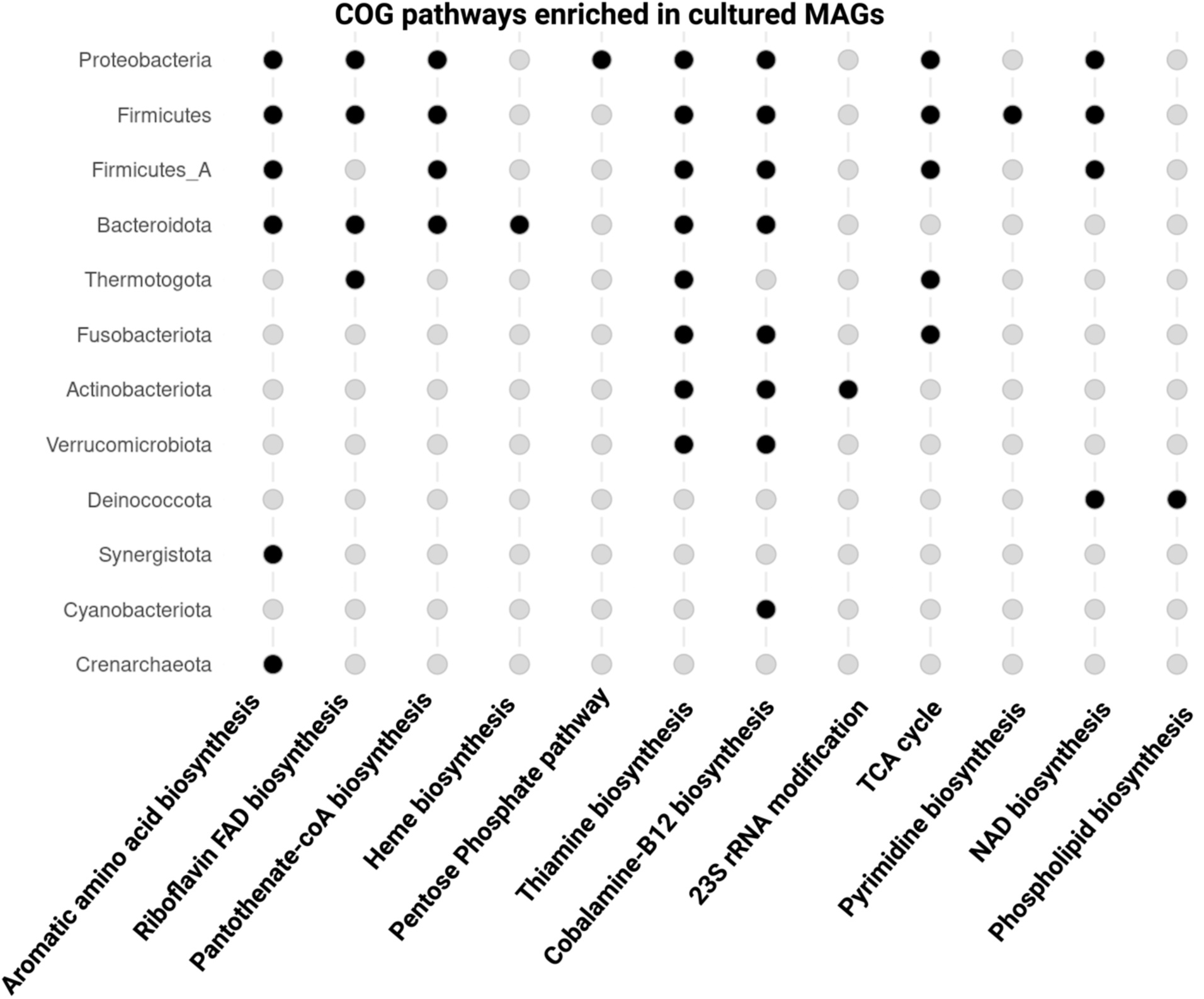
Clusters of Orthologous Groups (COG) pathways that are significantly enriched in cultured MAGs in at least one phylum and not enriched in uncultured MAGs in any phylum. Each column in the plot represents a COG pathway, and each row represents a phylum. Solid circles indicate phyla where the corresponding pathway is significantly enriched in cultured MAGs (Fisher’s exact test, FDR-adjusted P < 0.05).

**Fig 4a:**
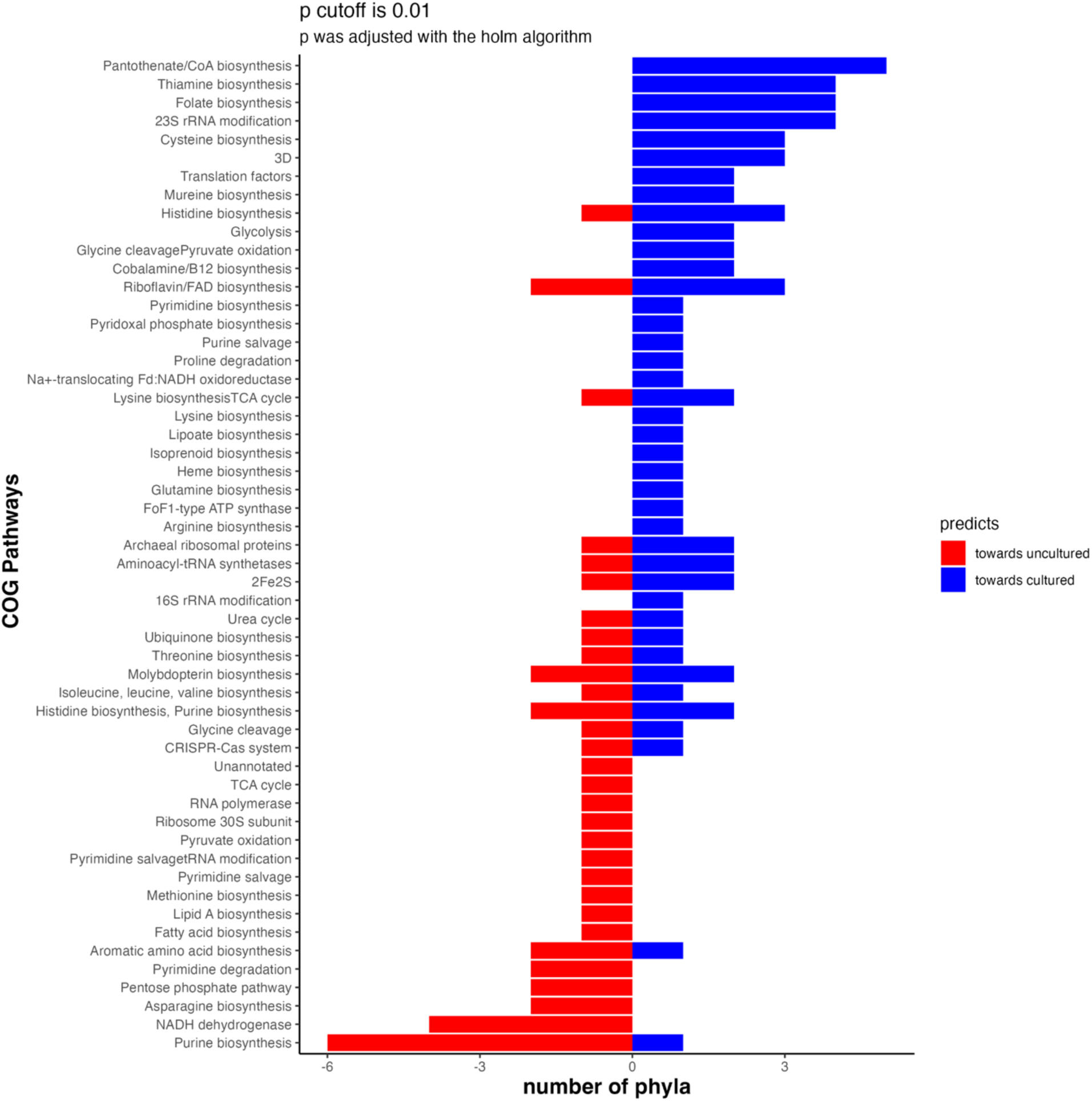
Machine learning (LASSO) predicts *important COG pathways that are predictive of microbial cultured or uncultured status. Pathways are filtered based on the permutation importance. Only pathways with permutation importance (predictive power > 0.1) are included in the plot*.

**Fig.4b:**
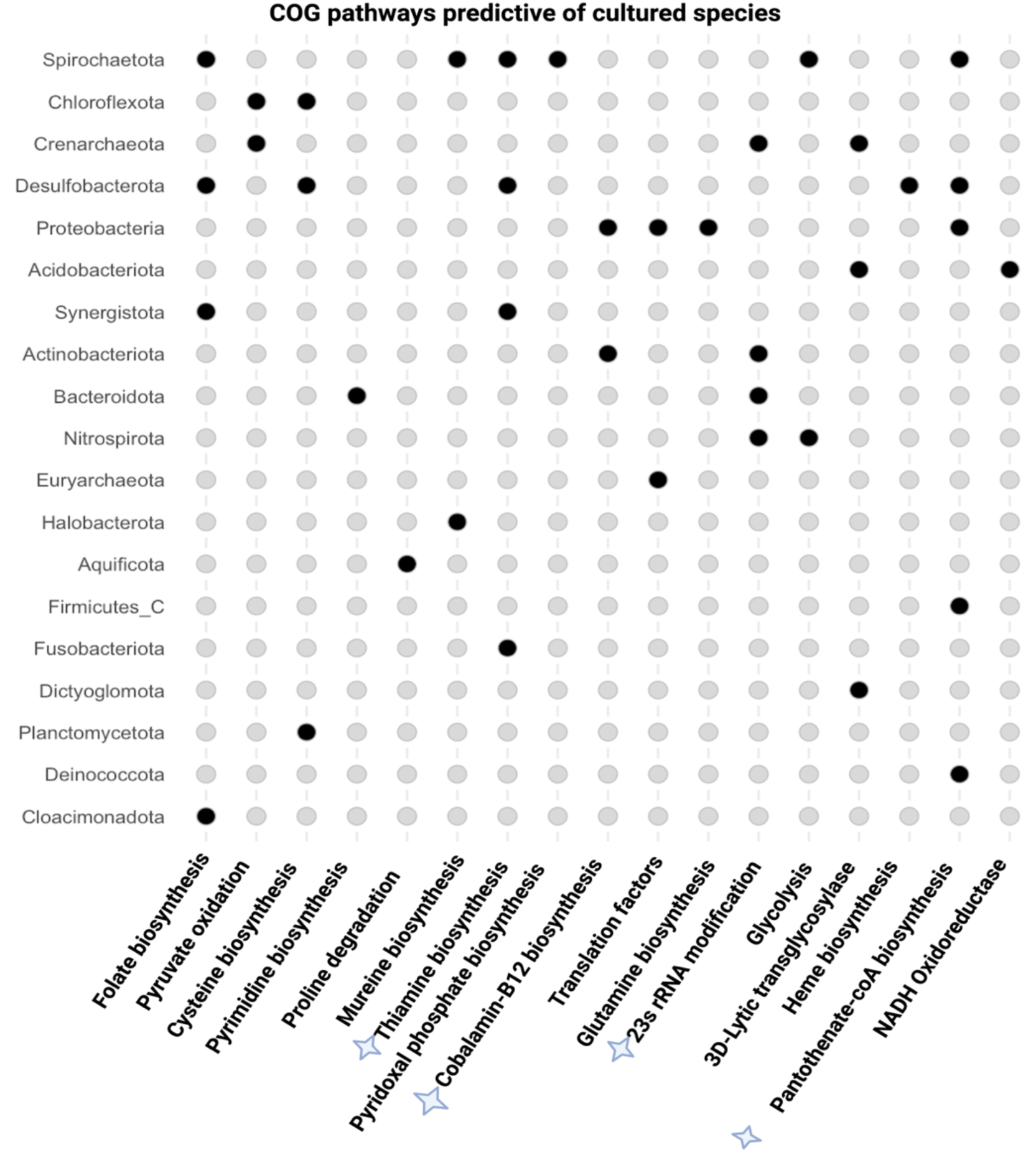
Machine learning (LASSO) predicts COG pathways that are strongly predictive of cultured status by phylum. Only COG pathways that predict cultured status with high permutation importance (>0.1) are included in the plot. Solid circles indicate the phyla for which this COG pathway is significantly predicts cultured status. COG pathways that are both enriched in cultured MAGs and predict microbial cultured status are starred and are predominantly vitamin biosynthetic pathways.

### LASSO regression identified metabolic predictors of culturability across phyla

To identify genomic features predictive of culturability, we applied LASSO regression to model culturability using the full set of COG pathway annotations. Unlike enrichment analysis, which tests each pathway independently, LASSO jointly considers all features while penalizing model complexity, producing sparse models that retain only the most informative predictors. We focused on pathways with nonzero LASSO coefficients and at least moderate permutation importance scores (>0.1), which reflect each pathway’s contribution to model performance. These pathways were identified in both well-sampled and poorly cultured phyla–including *Acidobacteriota*, *Planctomycetota*, *Spirochaetota*, and *Chloroflexota* (Fig. S4), suggesting that LASSO captures generalizable predictors of culturability across microbial diversity.

Consistent with the previous enrichment results, the strongest predictors of cultivability were dominated by cofactor and vitamin biosynthesis pathways. Thiamine, folate and pantothenate/CoA biosynthesis were frequently retained across phyla. For example, pantothenate/CoA biosynthesis predicted culturability in well-represented groups such as *Deinococcota*, *Proteobacteria*, *Spirochaetota*, *Desulfobacterota*, and *Firmicutes_C*. Thiamine and folate biosynthesis also predicted cultured status in multiple groups including *Spirochaetota*, *Desulfobacterota*, and *Synergistota*. In contrast, purine biosynthesis and NADH dehydrogenase were predictive of uncultured status and showed high permutation importance in multiple phyla, possibly reflecting metabolic dependencies or adaptations incompatible with growth in isolation.

### Culturability correlates with growth strategy and phylogeny

A heatmap of LASSO-derived feature importance across phyla (Fig. 5) revealed distinct clustering patterns that partially align with known microbial growth strategies. Fast-growing, frequently cultured phyla—such as *Bacteroidota*, *Firmicutes_C*, *Proteobacteria*, and *Actinobacteriota*—cluster together (Fig. 5, clade 2) and shared similar sets of culturability-associated pathways, suggesting conserved genomic predictors of laboratory growth.

**Fig 5:**
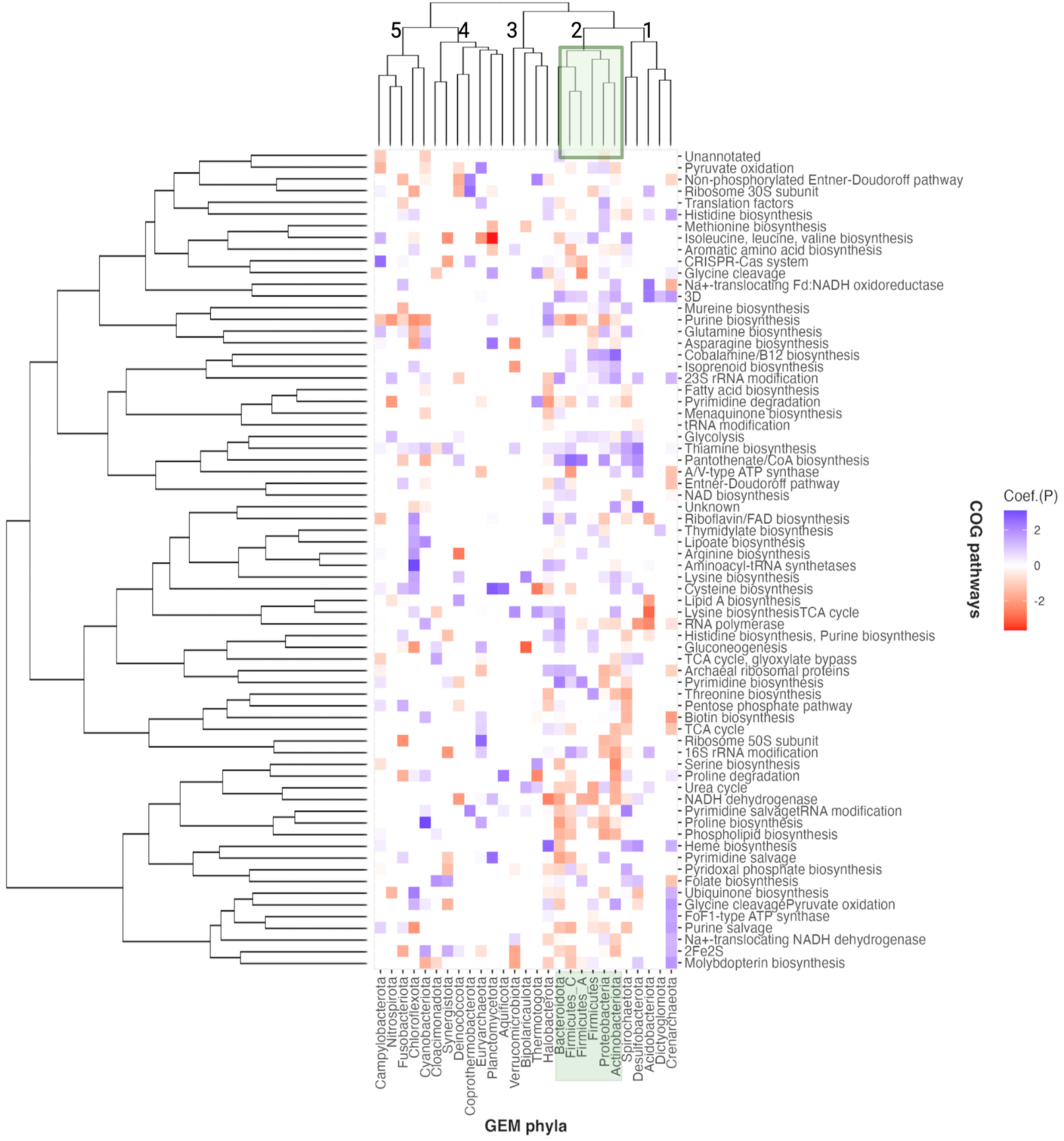
Heat map showing COG pathways predicting cultured (blue) and uncultured (red) species within each phylum. The intensity of the color represents the strength of the prediction (see methods). Various clusters are numbered and phyla in which fast growers are known to be abundant are highlighted in green. Microbes that are known to be fast growers and have higher cultured representatives cluster together while slow growers show greater variability in clustering.

In comparison, slow-growing or poorly cultured phyla such as *Spirocaetota*, *Desulfobacteriota*, *Acidobacteriota*, *Dictyoglomota*, and *Crenaracheaota* form a separate clade (clade 1) characterized by more variable or phylum-specific predictors. Strains from these phyla may require co-culture, longer incubations, oligotrophic media, or extremely long growth periods that are not routinely used in cultivation experiments.

Extremophilic phyla such as *Halobacteriota*, *Thermotogota*, and *Bipolaricaulota* form their own distinct cluster (Fig. 5, clade 3), suggesting that the genomic features associated with laboratory culturability in these lineages differ from those in mesophiles. This could reflect both biological traits and biases in culturing methods.

Finally, several phyla, including *Campylobacteriota*, *Nitrospirota*, *Chloroflexota*, exhibited mixed clustering patterns, containing both fast- and slow-growing lineages. This likely reflects a broader range of ecological niches and growth strategies within these phyla, resulting in more heterogeneous culturability-associated genomic features.

## Discussion

All microbes, including the best-studied model organisms, contain a substantial fraction of genes with unknown or poorly characterized functions. For example, even *E. coli* K-12 has ∼35% of genes lacking experimental annotation (22). In our analysis of over 52,000 metagenome-assembled genomes (MAGs) from the GEM catalog, we find that this “functional dark matter” (2, 23) is substantially more prevalent among uncultured microbes. After comparing uncultured MAGs to both cultured MAGs and isolate genomes, we find that uncultured taxa exhibit markedly greater predicted functional novelty (Fig. 1), as reflected in lower amino acid identity to SwissProt proteins and lower bitscores normalized by alignment length.

This novelty increases with genetic distance from cultured representatives: genes from MAGs that belong to cultured species show the highest similarity to characterized proteins, followed by those within the same genus, and so on (Fig. 2). This trend likely stems from the historical reliance on model organisms, selected in part for their ability to grow in pure culture, for protein function characterization (4, 13, 24).

Prior work has reported widespread genomic novelty among uncultured microbes from diverse environments (2, 15, 23, 25–27). Our results reinforce and extend this view, showing that uncharacterized genes are not random or artifactual, but are concentrated in lineages phylogenetically distant from cultured taxa and disproportionately prevalent in archaeal MAGs. The high proportion of genes with unknown function—especially in uncultured archaea— suggests extensive, unexplored functional diversity with likely biological significance (28–31).

To explore how genomic content relates to culturability, we examined COG pathway enrichment across 28 microbial phyla. Cultured MAGs were consistently enriched in pathways involved in cofactor and vitamin biosynthesis, energy metabolism (e.g., TCA cycle), and CRISPR-Cas defense (Fig. 3A). These cofactor and vitamin biosynthesis likely support autonomous growth in axenic lab conditions. TCA cycle and CRISPR Cas system are fundamental for rapid energy production, growth, and viral defense, potentially explaining their prevalence in highly cultured microbes. The TCA cycle can provide the energy efficiency to support the higher metabolic demands of fast-growing microbes (32, 33) while the CRISPR-Cas system can help microbes fend off viral infections, especially in environments where exposure to phages could limit their survival (34, 35). In contrast, uncultured MAGs were enriched for pathways such as NADH dehydrogenase activity, iron-sulfur cluster biosynthesis, and asparagine metabolism—features potentially associated with ecological dependencies or niche-specific adaptations (36–39).

Notably, we identified 15 pathways that were exclusively enriched in cultured MAGs and absent from uncultured ones (Fig. 3B). These include biosynthetic pathways for thiamine, folate, pantothenate/CoA, and cobalamin, which are well-documented microbial growth factors (11, 40, 41). Our findings support the hypothesis that biosynthetic self-sufficiency is a strong predictor of culturability. For instance, Zheng et al identified uncultured *Burkholderiales* genomes from sludge samples as dependent on vitamin B12 (cobalamin) and their experimental findings revealed that the addition of vitamin B12 stimulated the growth of these taxa (42).

Vitamins are among the most frequently exchanged metabolites in microbial communities (43), and microbes often rely on co-produced vitamins for growth (44, 45). For example, *Variovorax* species support thiamine auxotrophs in bioreactor microbial communities while depending on pantothenate supplied by other species (46). Such interdependencies—often mediated by unidentified small molecules—are difficult to replicate in vitro (47, 48). Facilitating small-molecule exchange via co-culture systems or in situ cultivation platforms (e.g., iChip) has significantly improved culturability (7, 10, 49). These findings suggest that many uncultured microbes rely on metabolite sharing, limiting their growth in isolation.

To identify genomic features that generalize across phyla, we applied LASSO regression to model culturability. LASSO consistently identified cofactor biosynthesis—particularly thiamine, folate, and pantothenate—as predictive of culturability, even in poorly sampled or rarely cultured phyla such as *Acidobacteriota*, *Planctomycetota*, and *Spirochaetota* (Fig. S4). In contrast, NADH dehydrogenase and purine biosynthesis were retained as predictors of uncultured status across multiple phyla (Fig.4a). Conversely, purine biosynthesis emerged in our analyses as a strong genomic predictor of uncultured status across multiple phyla (Fig. 3). Because purine biosynthesis is metabolically expensive (50, 51), microbes that invest in it may be outcompeted in nutrient-rich laboratory media where purines are abundant. This suggests that uncultured microbes may favor strategies incompatible with current cultivation protocols. Together, these analyses highlight overlapping but distinct predictors of culturability.

Clustering of phyla based on LASSO-derived predictors further revealed meaningful culturability patterns (Fig. 5). Fast-growing phyla that have many cultured members, such as *Bacteroidetes* and *Firmicutes*, clustered tightly and shared a relatively uniform set of culturability-associated pathways. By contrast, slow-growing phyla, including *Spirochaetota*, *Desulfobacterota*, and *Acidobacteriota*, clustered separately and displayed greater pathway heterogeneity. These patterns likely reflect a broader range of growth strategies, ecological adaptations, and unrecognized metabolic dependencies. This also suggests that culturability-linked features are more conserved among fast-growing taxa.

Taken together, our results indicate that culturability is a complex, multifactorial trait shaped by biosynthetic independence, energy metabolism, and ecological interdependence. Cofactor biosynthesis emerges as a consistent feature of culturability, while pathways associated with ecological dependency or high energetic cost (e.g., purine biosynthesis) are more common among uncultured lineages.

These insights challenge the notion that uncultured microbes remain so due mainly to oversight or insufficient effort. Instead, they highlight fundamental differences in metabolic and ecological requirements from the microbes themselves. Targeted, genome-informed cultivation, especially approaches that supplement key vitamins or simulate inter-species interactions, offers a promising roadmap for accessing the hidden functional potential of uncultured microbes.

In conclusion, our study provides robust evidence that uncultured microbes encode greater predicted functional novelty and contain distinctive genomic predictors of culturability. Biotechnological discovery may be enhanced by concentrating on uncultured clades, especially among the archaea. By identifying generalizable pathways associated with cultured status, we offer a framework for prioritizing taxa and traits for experimental cultivation. Unlocking these microbes will not only expand the known tree of life, but also reveal the vast and largely unexplored landscape of microbial functional diversity.

## Methods

### Description of data sets

#### GEM

The GEM (Genomes from Earth’s Microbiomes) data comes from the GEM catalog, which consists of 52, 505 metagenome-assembled genomes (MAGs) assembled from a diverse set of environments sampled from around the world including continental soils, oceans, agricultural soils, animal and human hosts. Of these, about 17.4% of the total MAGs were classified as high-quality (≤5% contamination ≥90% completeness, ≥18/20 rRNA and tRNA genes), and the remaining MAGs were medium quality according to the minimum information about a Metagenome Assembled Genome (MIMAG) standard. Hence, each MAG has a completeness level greater than 50% and a contamination level less than 5% as estimated with CheckM (v1.0.11). The GEM data have been classified taxonomically at higher ranks using GTDB and clustered on 95% ANI to derive the species-level Operational Taxonomic Unit (OTU). The species-level OTUs assignment of these MAGs was cross-validated with that of the GTDB taxonomy (15). GEM MAGs were annotated for Cluster of Orthologous Genes (COGs) pathways and genes in the anvio environment using the “anvi-run-ncbi-cogs” commands

#### Isolates

The isolates dataset comprises 7,035 genomes sourced from the IMG/JGI database, downloaded prior to 2020, and filtered to include only cultured isolates.

### Cultured Status

To infer the cultured status of each GEM MAG, we first compared taxonomic annotations of each MAG to those RefSeq category (excluding any uncultured genomes such as single amplified genomes or metagenome-assembled genomes) of the same GTDB release (vs 89). MAGs were classified to the lowest taxonomic level at which they have a cultured representative, for instance if a MAG matched a RefSeq genome taxonomically ranked to an order level then the MAG has the corresponding cultured level. b) We next aligned the sequences of each MAG in the GEM dataset to NCBI isolate RefSeq genomes via Reference Seeker (52) to find if there are any closely related reference genomes. Reference Seeker first conducted a Kmer-based genome distance between the GEM MAGs and RefSeq with Mash (53) and then calculated the Average Nucleotide Identity (ANI) to determine the MAGs with ANI > 95% and conserved DNA > 69%, a commonly used threshold for delineating the prokaryotic species level. MAGs with species cultured representative or matched an isolate RefSeq with ANI > 95% are termed as “cultured MAGs” while MAGs that do not meet this criterion are referred to as “uncultured MAGs”

### Comparing The Genes in Cultured Genomes to Uncultured Genomes

To determine the similarity of sequences from cultured and uncultured genomes to SwissProt, a database whose gene products have been rigorously characterized. We conducted alignments between each gene in the GEM dataset, translated into amino acid sequences, and to the genes in the SwissProt dataset using diamond’s "blastp" command (54).

Diamond’s “blastp” command was also conducted on the cultured isolates group. We set the e-value cutoff to 100 instead of the default e-value because we are interested in any slightest similarity of the GEM sequences to Swissprot. This cutoff ensures many false positive alignments and that even minor similarities between GEM genes and SwissProt genes are identified. We deal with false positives separately by comparing the sequence similarity of random sequences simulated from the GEM database to the SwissProt. The BlastP reports were excluded to only the top hits of the query search. We found that 112,784,914 out of 122,978,191 protein sequences in the GEM MAGs aligned to genes in the SwissProt database, and 27,952,866 out of 30,840,483 protein sequences of the isolates aligned to the SwissProt database sequences. The bit scores were normalized by dividing them by the alignment length. We also generated a random seq from the GEM MAGs dataset using the same e-value. We further compared the sequences in the GEM MAGs at each taxonomic cultured level to the Agnostos HMM profile database to quantify the genes with known and unknown functions (Supplemental Figure 2.3).

### Fisher’s enrichment test analysis

To identify the functional pathways and genes that could provide insights into potential factors contributing to cultivability, we analyzed the COGs enriched in cultured or uncultured MAGs. For each phylum, we did comparative statistical analysis of COGs presence or absence using Fisher’s exact test and enriched COGs in either cultured MAGs or uncultured MAGs based on relative proportions. We specifically focused on COG pathways that are significantly over-represented or under-represented in each group (P < 0.05).

### Description of the LASSO regression model pipeline for analyzing cultured status

To assess the distinction between cultured and uncultured GEM microbes, we employed LASSO regression to predict if an annotated pathway can be associated with cultured status. We designed a machine learning-based pipeline specifically intended to predict metabolic pathways associated with cultured status while mitigating the effects of phylogeny. To create the custom dataset, we assigned a label of “cultured” to the cultured MAGs or “uncultured” if it belonged to any other taxonomic cultured level. The dataset contained pathway features, which represent coarse-grained COG assignments (aggregated into higher biochemical pathways). The value of each feature is the approximate number of pathways in a given genome/species with a specific COG assignment. We normalized these pathways by abundance counts to accurately represent the phylum. To partially control for phylogeny, we remove all metadata features from the feature set used in the model, as we have previously shown metadata to be overly predictive of culturability regarding this particular dataset (55). These metadata include the cultured levels, taxonomy classes and taxonomic distances, completeness, contamination, and genome size. To finish controlling for phylogeny, we subset the data by phylum and run the machine learning part of our pipeline separately on each phylum (Fig. S5), effectively controlling for taxonomy. The machine learning section of our pipeline consists of the following steps: (i) preprocess the data, (ii) train a LASSO-based bagging regressor to predict cultured or uncultured status, and (iii) compute LASSO coefficient confidence intervals and feature importances. The details of these steps can be found in the supplementary materials.

## Data availability

The GEM MAGs and metadata are publicly available at https://portal.nersc.gov/GEM or https://genome.jgi.doe.gov/GEMs and the isolates are publicly available at https://img.jgi.doe.gov/

## Code availability

The code describing each of these steps can be found on GitHub (https://github.com/ababjac/GEM-feature-selection)

## Acknowledgments

The authors acknowledge funding from the National Science Foundation (OCE-2145434) and the Genomic Science Program of the U.S. Department of Energy, Office of Science, Office of Biological and Environmental Research (DE-SC0020369 to A.D.S. and K.G.L.). Additional funding came from the National Science Foundation (OCE-2152551 and EAR-2121637) to K.G.L. Further, A.N.B was supported by the National Science Foundation GRFP (Graduate Research Fellowship Program) during the completion of this work.

## SUPPLEMENTAL METHODOLOGY

### Classifying GEM data set into taxonomic cultured levels

Although in theory any species can be assigned a taxonomy to the species level, in practice, microbes with no close relation in the Genome Taxonomy Database (GTDB) will not be assigned to lower specific taxonomic ranks (species, genus…). Thus, the level of assigned taxonomic rank is a proxy for the frequency of an organism’s close relatives in GTDB. A larger fraction of bacterial GEM MAGs was assigned to lower taxonomic ranks and more frequently to taxa with cultured representatives than Archaea, which suggests that bacteria tend to be more cultured (Fig S2). For instance, 52.4% of bacteria GEM MAGs were assigned to a species vs. 43.3% for archaeal MAGs. 22.2% of bacterial MAGs were classified as a species with a cultured representative, whereas only 10.5% of archaeal MAGs were classified as the same species-cultured group (Fig. S2b). At the other end of the taxonomic rank spectrum, 47.6% of bacterial GEM MAGs were taxonomically ranked only to the genus level while for the archaeal MAGs, 56.7% were taxonomically ranked only to the genus level. (Fig S2a). For the taxonomic cultured groups, 73.7% of Bacterial MAGs were taxonomically classified up to the same genus level as a cultured organism, whereas 78.7% of archaeal MAGs were classified as being cultured (Fig. S2b.). 4.1% of the bacterial MAGs were not taxonomically classified to any cultured organism, which is a lesser percentage than the 10.8% proportion of the archaeal MAGs. Since GTDB includes sequences from both cultured and uncultured organisms for taxonomic assignment, this indicates that a reasonable percentage of the microbial taxonomic information was derived from uncultured groups.

### Details of necessary machine learning preprocessing

The preprocessing consists of standard machine learning practices: (i) stratify the data into training/testing sets, (ii) scale the data to meet model requirements, and (iii) rebalance by phylum to create a balanced training set. We split the data using the train_test_split function from scikit-learn, with 20% reserved for validation, and 80% utilized for training and stratification performed based on the cultured labels. We scale the data using the MinMax normalization technique with standard parameters. We then rebalance using SMOTE (1) which is an oversampling technique that creates new samples in a feature space by selecting along the path of two closely related real data points. We additionally note that any phyla that does not have a minimum of ten samples total and/or has less than three samples of either label been removed from the analysis (Table S2); this is due to statistical constraints on the number of samples that are required to use the methodologies presented here.

### Details of LASSO-based regression model for predicting cultured and uncultured status

We employed LASSO (Least Absolute Shrinkage and Selection Operator) as the regression component of our analysis pipeline as it has been previously shown to be a robust feature selection tool (2)and to be a comparable dimensionality reduction method to neural network-based architectures on these data (3). LASSO is a modified form of regression that uses a cost function to tune the model based on an *α* hyperparameter (4). The lower the *α* parameter, the more LASSO begins to approximate standard regression, whereas higher *α* values tend to cause coefficients to more rapidly trend toward zero. When using LASSO, features are selected based on the coefficient, i.e., if the coefficient is non-zero despite the stringent *α* penalty, then we say it contributes to the model.

We incorporated LASSO as the underlying model in a bagged regressor -- an ensemble model that is the composite of fitting a base model (LASSO) on many random subsets of the data. We build our bagging regressor using scikit-learn’s Bagging Regressor with estimators set to fifty and bootstrapping set to True. Each estimator in the bagged regressor is an instance of a logistic regression (scikit-learn’s Logistic Regression) model where the penalty is set to “l1” representing a LASSO penalty. We train a separately bagged regressor for each phylum using 5-fold cross-validation within the 80% training partition (Fig.S5).

### Details regarding calculating and interpreting coefficients, confidence intervals and permutation importance

A bagged regressor is advantageous because it mitigates the effect of randomness as LASSO (like many other machine learning methods) will return slightly different features based on the random seeding. This not only makes our predictions more verbose but also allows for calculating t-confidence intervals for each coefficient. For each feature, we calculate the mean and standard deviation of its associated coefficients from the underlying LASSO models in the trained bagged regressor. We then compute standard confidence intervals and p-values using the t-distribution. Confidence intervals and p-values are directly linked in that smaller p-values will have larger confidence intervals because we are more confident, that we contain the true value. Since we are looking for non-zero coefficients based on LASSO, we can select a feature if its confidence interval does not contain zero. We tested two cutoffs for selecting significant coefficients based on the p-values (p < 0.05 and p < 0.1) and found that p < 0.05 had a stronger signal. If a coefficient is significant, we then interpret it as follows: “For each (normalized) 1 unit increase of a sample’s feature_X, we would expect an average coefficient_X increase/decrease in the probability that sample is cultured”, where increase/decrease comes from the value of the coefficient.

A common misconception when interpreting coefficients of regression models is that larger coefficients are more “important” to model understanding. To determine the most important coefficients we additionally perform permutation importance testing (5) using the 20% validation data which was held out from the original modeling. The permutation importance algorithm is an explainable AI method that calculates how much each feature contributes to a trained model by permuting over the fitted estimator (our LASSO bagged regressor in this case) and randomly shuffling one feature each time. When this feature is removed, we can get a measure of what the model is “missing” by randomizing it. Higher permutation importance scores indicate an increasingly positive influence on the model, while scores below zero suggest the feature may have negatively affected model performance. We then put this feature back into the model and remove the next, iterating through all features to get a full set of importances. By doing this on the validation data, we get a measure of each feature’s generalizability to new data. We used Scikit-Learn’s permutation importance algorithm for these calculations (6). This algorithm was chosen because it should be more robust to feature inflation than impurity-based feature selection methods (e.g. Gini).

### Shuffling data to create a random baseline

As an additional sanity check, we aim to ensure the performance and coefficients from our analysis cannot occur by random chance. A standard technique for validating experimental results is to perform shuffling of the input data before repeating the analysis and comparing the results to those of the true data. We create a shuffled dataset using Scikit Learn’s shuffle algorithm with default parameters. We omit both the cultured labels and metadata columns from being shuffled to preserve the integrity of the phyla and predictions when modeling. We permute five different partitions of shuffled data and average the results to produce our random baseline set. We replicate all steps of the analysis using the shuffled data as described previously.

### Evaluating model performance

In order to evaluate our models, we use standard classification metrics: AUROC, accuracy, precision and recall. AUROC represents the area under the receiver operator curve which is a number between 0 and 1 showing a true vs false positive rate across all classification thresholds (7). For balanced data, we would expect a minimum AUROC of 0.5 if we predicted everything as true for a baseline comparison. Accuracy measures the proportion of correct predictions with respect to the total number of predictions (i.e., Accuracy = TP + TN / Total). Precision accounts for the proportion of correct positive identifications (i.e., Precision = TP / TP + FP). Finally, Recall identifies the proportion of correctly classified positive labels (i.e., Recall = TP / TP + FN). We compute each metric using the 20% validation set per phylum. We determine a minimum threshold for “good” performance based on the shuffled data, i.e., if the real performance is statistically greater than the shuffled performance, we say that the model is performing better than we would expect by random chance. We also note that this dataset (GEM) was previously baselined against several other methods (3) but not by individual phylum.

To confirm our LASSO performance is better than what would occur by random chance, we repeated our analysis (see Fig. S5) on shuffled data, i.e., data for which the null hypothesis of no genes or gene clusters being associated with culturability was known to be true (see Methods). Figure S6 shows the performance comparison for evaluating the real data (orange) and shuffled data (blue), where the real data outperforms the shuffled data on average across all the performance metrics considered. Accuracy and AUROC are both greater than 0.8 for most phyla, whereas shuffled performance sits around 0.4-0.7 and 0.4-0.6 respectively (Fig. S6). The Recall is similarly greater than 0.8, and the Precision is greater than 0.7 for nearly all phyla. The difference is more striking with Precision and Recall as most shuffled phyla are well below a score of 0.5 for these metrics which suggests a high level of misprediction in our random baseline that does not occur in our real models. There are only a few phyla where the shuffled performance was within a margin of error of the real performance based on Accuracy (Acidobacteriodata, Coprothermobacterota, and Planctomycetata) (Fig.S6). After further inspection, these phyla are highly unbalanced with few cultured/uncultured representatives (Table S2) which explains the similar Accuracy for both real and shuffled data. When looking at other metrics for these phyla that better handle imbalance (AUROC, precision, recall), we see that these phyla do outperform random chance in terms of false positives/negatives. We conclude based on these results that the performance of all phyla could not occur by random chance. This confirms that our LASSO models are capturing biologically important signals from both the pathway and annotation features that can be interpreted in the next sections.

## SUPPLEMENTAL FIGURES

**Table S1:**
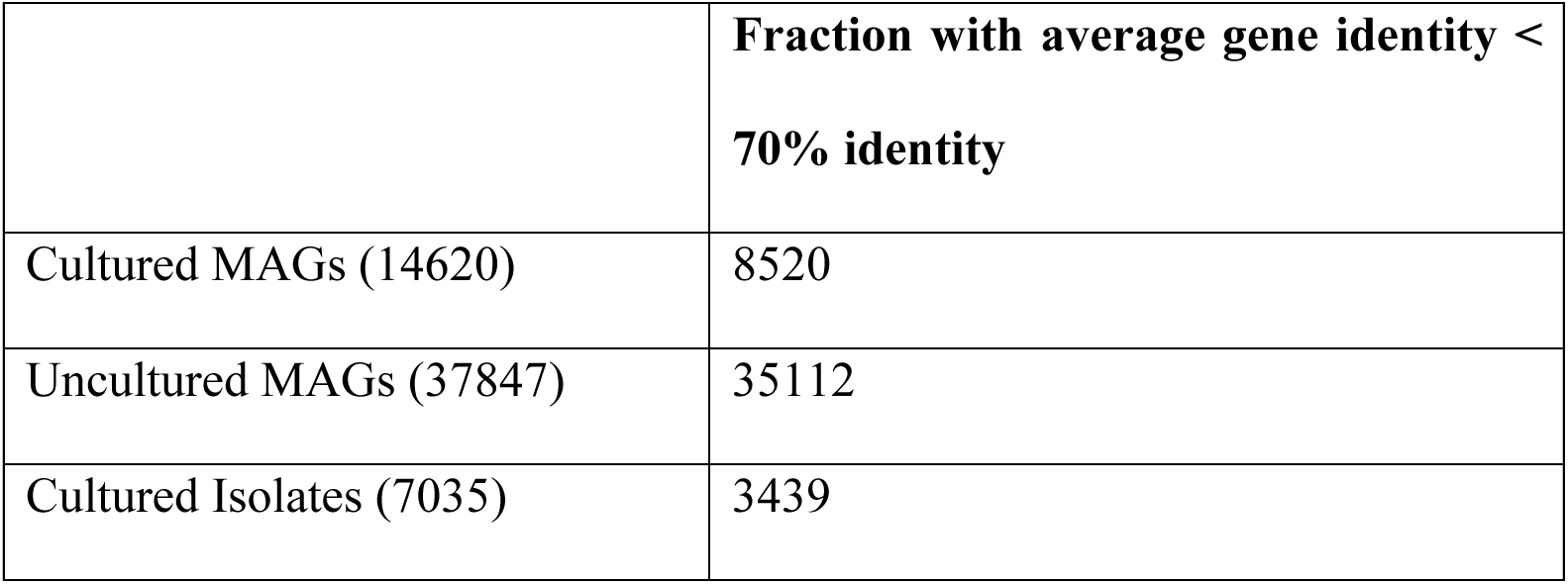
Counts of cultured and uncultured MAGs with average gene identity < 70% to SwissProt.

|  | Fraction with average gene identity < 70% identity |
| --- | --- |
| Cultured MAGs (14620) | 8520 |
| Uncultured MAGs (37847) | 35112 |
| Cultured Isolates (7035) | 3439 |

**Table S2:** Counts of cultured and uncultured samples per phylum based on the taxonomy of cultured species used in LASSO regression analysis. Phyla are excluded from analysis (and therefore also from this table) if either category contains less than 5 samples (see Methods).

| Phylum | Cultured Counts | Uncultured Counts |
| --- | --- | --- |
| Fusobacteriota | 81 | 134 |
| Planctomycetota | 8 | 894 |
| Deinococcota | 145 | 55 |
| Firmicutes_A | 1962 | 6853 |
| Cyanobacteriota | 10 | 478 |
| Firmicutes | 658 | 1301 |
| Firmicutes_C | 599 | 654 |
| Coprothermobacterota | 10 | 6 |
| Chloroflexota | 40 | 1112 |
| Verrucomicrobiota | 208 | 1202 |
| Thermotogota | 25 | 176 |
| Acidobacteriota | 5 | 739 |
| Proteobacteria | 2190 | 8460 |
| Cloacimonadota | 11 | 107 |
| Campylobacterota | 85 | 279 |
| Bacteroidota | 2776 | 6266 |
| Actinobacteriota | 841 | 3210 |
| Desulfobacterota | 72 | 910 |
| Aquificota | 9 | 44 |
| Synergistota | 18 | 135 |
| Nitrospirota | 16 | 153 |

**Figure S1:**
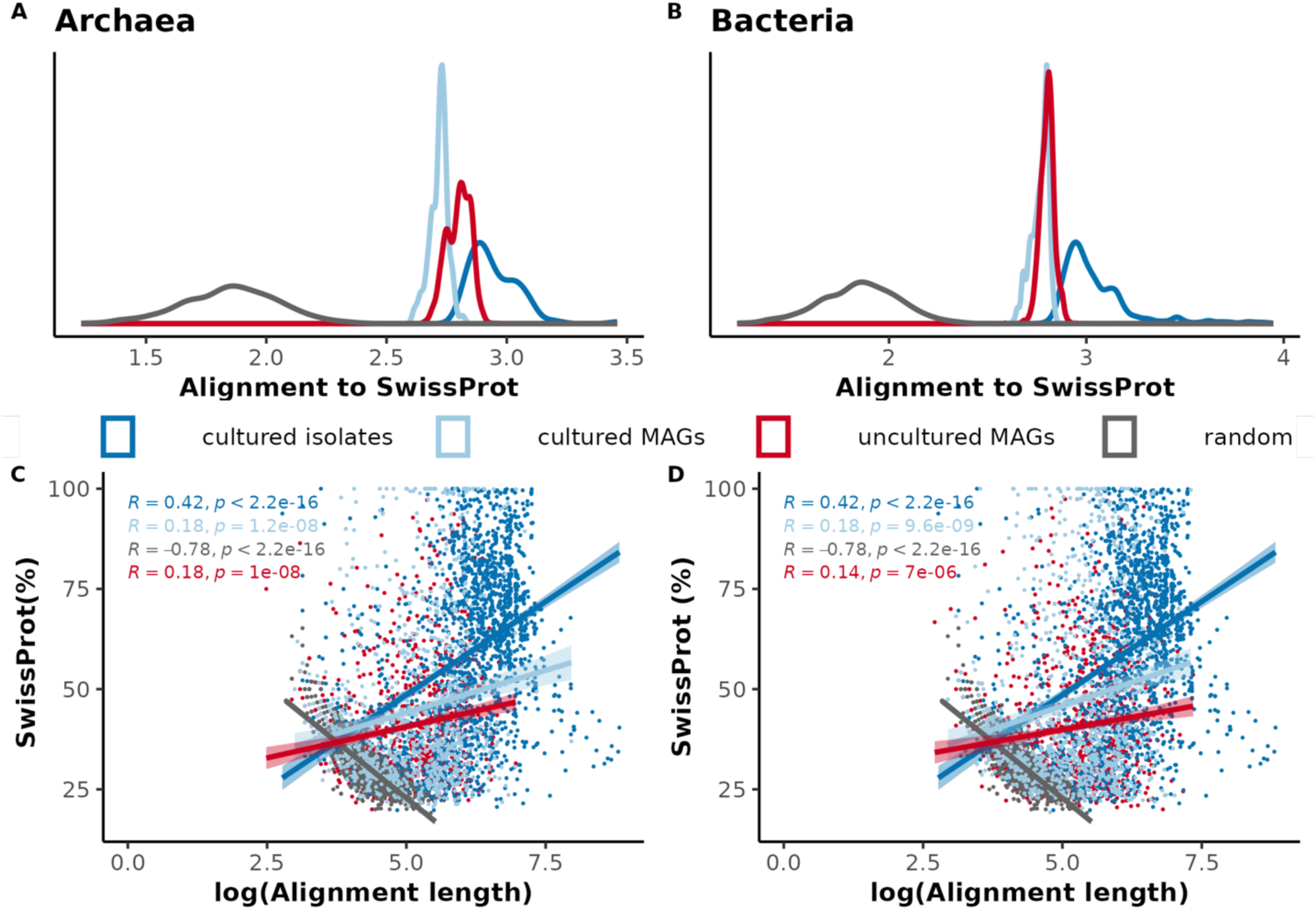
Distribution of mean of alignment length of MAGs, isolates, and random sequences to SwissProt in A) Archaea and in B) Bacteria. C and D represent the relationship between alignment length and identity to SwissProt of genes (1000 samples) from cultured isolates, uncultured MAGs, cultured MAGs Archaea and Bacteria respectively

**Figure S2:**
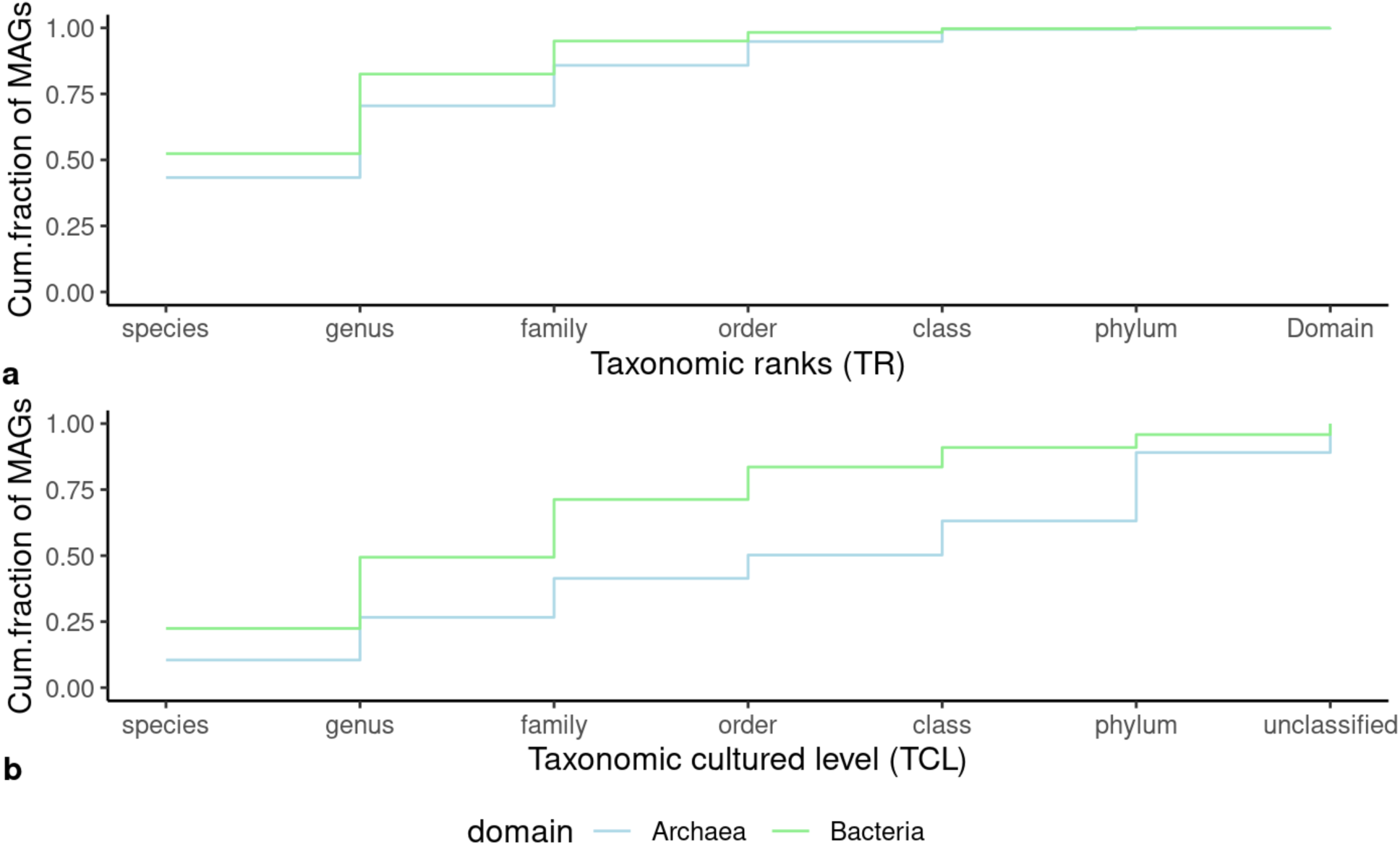
Cumulative fractions of taxonomic cultured representatives in GEM data sets. The GEM MAG lowest taxonomic rank was assigned by Nayfach et al. (2021) using all Genome Taxonomy Database (GTDB) reference genomes (Chaumeil et al., 2020) while the taxonomic cultured levels were assigned based on the RefSeq category of the same GTDB release excluding single-cell amplified genomes and the assembled metagenomes.

**Figure S3:**
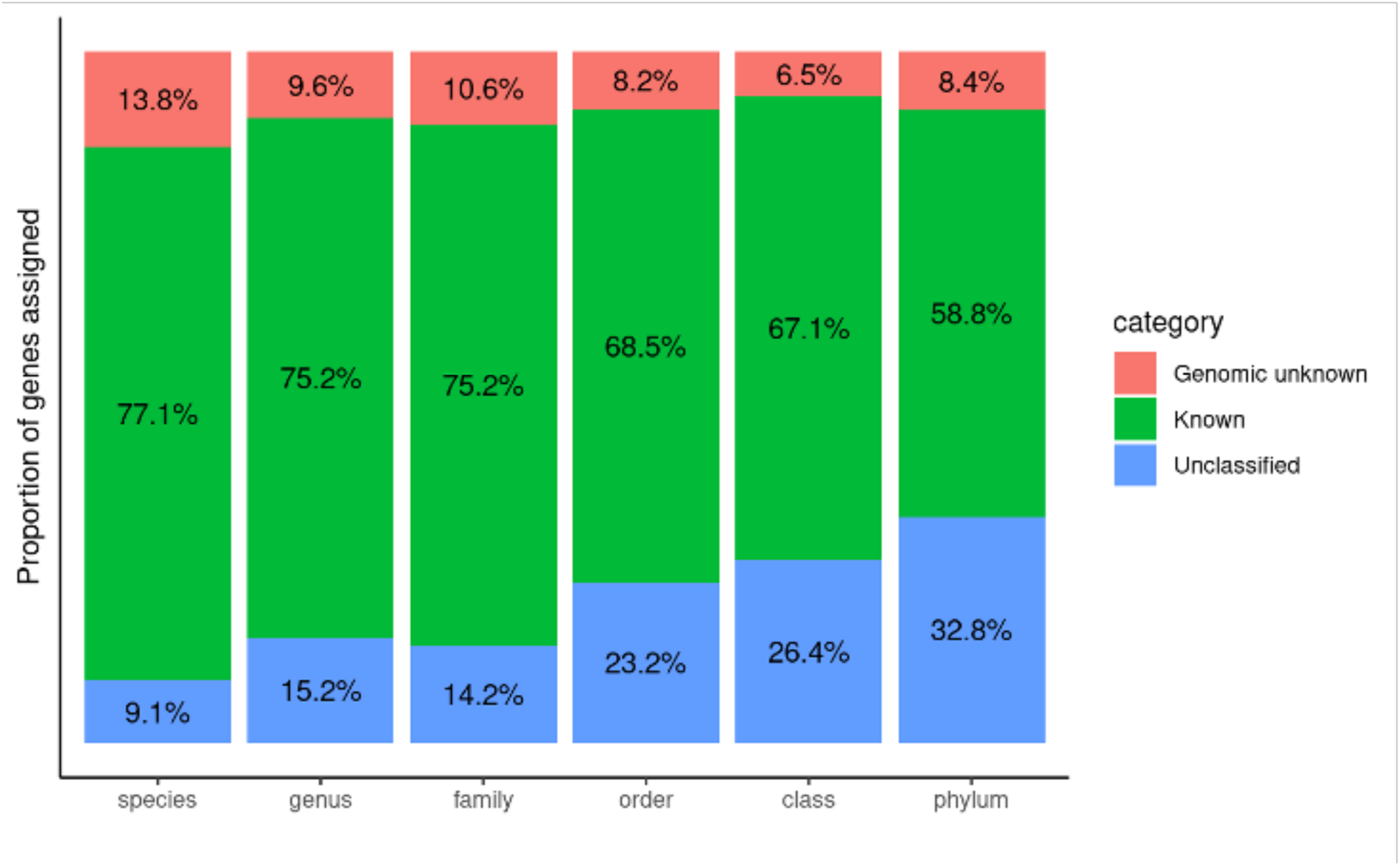
Proportion of genes assigned by the Agnostos DB clusters profile search database at different GEM taxonomic cultured levels.

**Figure S4:**
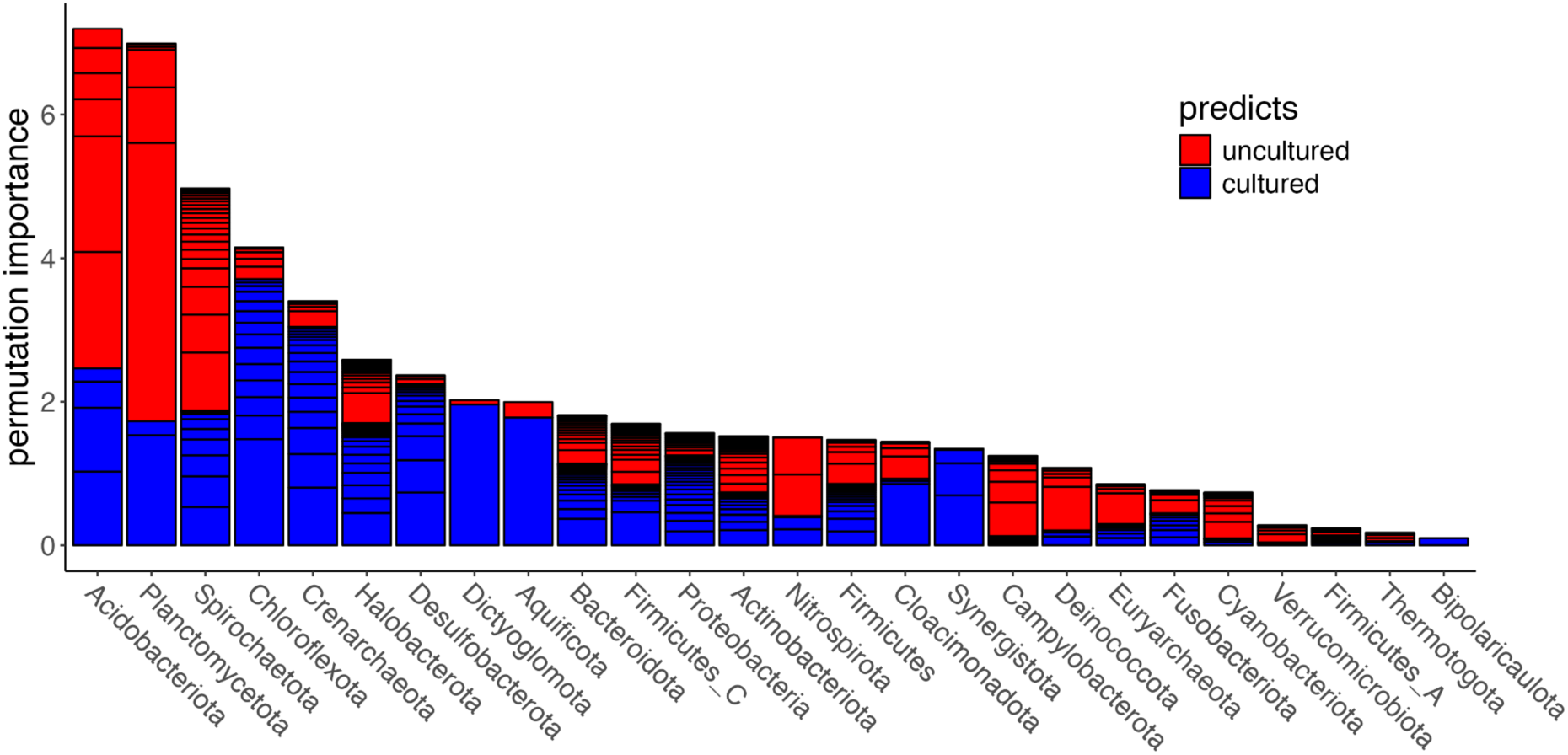
Permutation importance by GEM phyla. Pathways from less cultured phyla tend to have COG pathways with higher permutation importance.

**Figure S5:**
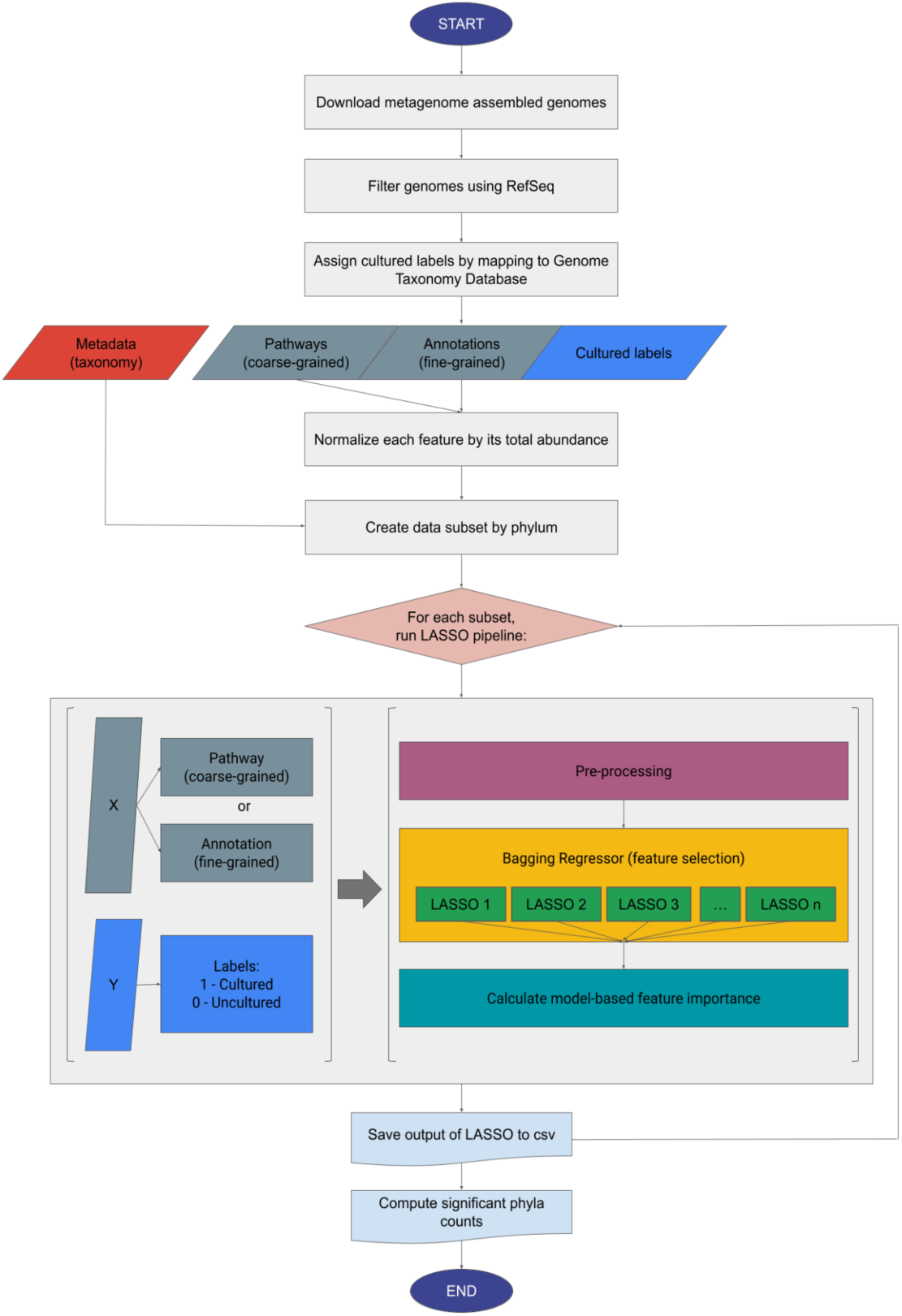
A flowchart showing our full methodology from start to finish. We begin by gathering metagenome information from the Genomes from Earth’s Microbiomes (GEM) project. We then create a custom dataset containing metadata, pathway features, annotation features, and cultured labels (See Methods). Each of these data are then split into subsets representing their ndividual phyla and the metadata are removed from the subsets. We remove any subsets that do not meet the minimum data requirements (see Methods). We lastly run our LASSO pipeline 5 times on each dataset and agglomerate the counts of significant phyla (as determined by confidence intervals (see Fig. S5d)) into a final output file to be analyzed.

**Figure S6:**
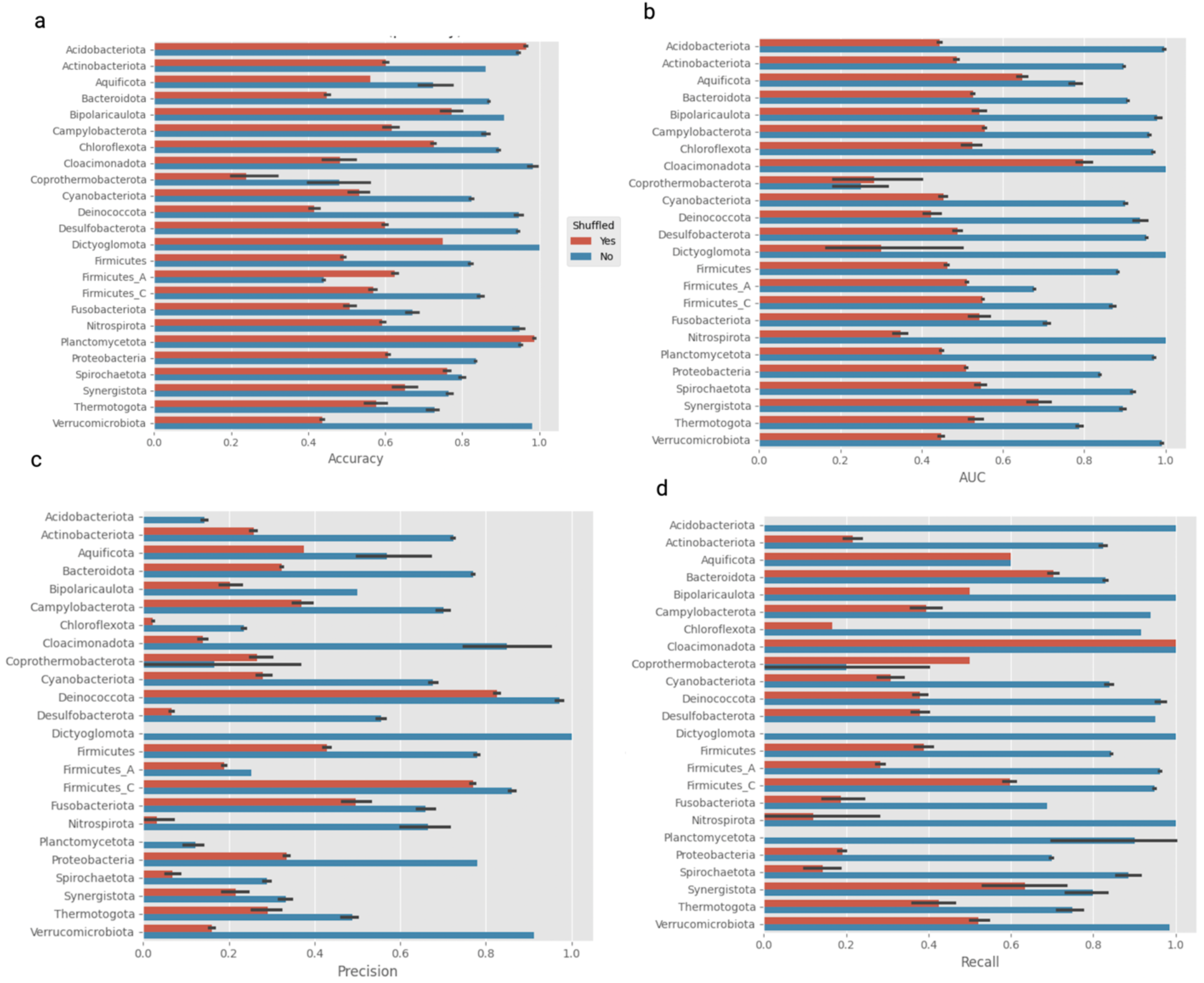
Performance metrics for GEM annotations and pathways: A) accuracy, B) AUC, C) Precision, and D) Recall (see Methods). For Accuracy, Precision and Recall a value of 0.0 is the worst and 1.0 is the best, whereas for AUC the worst value is actually 0.5. The orange represents real data (i.e. shuffled=”No”) and the blue represents our random baseline (i.e. shuffled=”Yes”). Phyla are omitted from interpretation if the margin of error of shuffled performance is within our actual predictions across all metrics.

## References

1. Lloyd KG, Steen AD, Ladau J, Yin J, Crosby L. 2018. Phylogenetically novel uncultured microbial cells dominate earth microbiomes. MSystems 3:e00055–18.

2. Rinke C, Schwientek P, Sczyrba A, Ivanova NN, Anderson IJ, Cheng J-F, Darling A, Malfatti S, Swan BK, Gies EA. 2013. Insights into the phylogeny and coding potential of microbial dark matter. Nature 499:431–437.

3. Steen AD, Crits-Christoph A, Carini P, DeAngelis KM, Fierer N, Lloyd KG, Cameron Thrash J. 2019. High proportions of bacteria and archaea across most biomes remain uncultured. ISME J 13:3126–3130.

4. Lewis WH, Tahon G, Geesink P, Sousa DZ, Ettema TJ. 2021. Innovations to culturing the uncultured microbial majority. Nat Rev Microbiol 19:225–240.

5. Mu D-S, Ouyang Y, Chen G-J, Du Z-J. 2021. Strategies for culturing active/dormant marine microbes. Mar Life Sci Technol 3:121–131.

6. Vartoukian SR, Palmer RM, Wade WG. 2010. Strategies for culture of ‘unculturable’bacteria. FEMS Microbiol Lett 309:1–7.

7. Berdy B, Spoering AL, Ling LL, Epstein SS. 2017. In situ cultivation of previously uncultivable microorganisms using the ichip. Nat Protoc 12:2232–2242.

8. D’Onofrio A, Crawford JM, Stewart EJ, Witt K, Gavrish E, Epstein S, Clardy J, Lewis K. 2010. Siderophores from neighboring organisms promote the growth of uncultured bacteria. Chem Biol 17:254–264.

9. Ghoul M, Mitri S. 2016. The ecology and evolution of microbial competition. Trends Microbiol 24:833–845.

10. Imachi H, Nobu MK, Nakahara N, Morono Y, Ogawara M, Takaki Y, Takano Y, Uematsu K, Ikuta T, Ito M. 2020. Isolation of an archaeon at the prokaryote–eukaryote interface. Nature 577:519–525.

11. Kamagata Y, Tamaki H. 2005. Cultivation of uncultured fastidious microbes. Microbes Environ 20:85–91.

12. Prakash O, Shouche Y, Jangid K, Kostka JE. 2013. Microbial cultivation and the role of microbial resource centers in the omics era. Appl Microbiol Biotechnol 97:51–62.

13. Garza DR, Dutilh BE. 2015. From cultured to uncultured genome sequences: metagenomics and modeling microbial ecosystems. Cell Mol Life Sci 72:4287–4308.

14. Handelsman J. 2004. Metagenomics: Application of Genomics to Uncultured Microorganisms. Microbiol Mol Biol Rev 68:669–685.

15. Nayfach S, Roux S, Seshadri R, Udwary D, Varghese N, Schulz F, Wu D, Paez-Espino D, Chen I-M, Huntemann M. 2021. A genomic catalog of Earth’s microbiomes. Nat Biotechnol 39:499–509.

16. Dunbrack Jr RL. 2006. Sequence comparison and protein structure prediction. Curr Opin Struct Biol 16:374–384.

17. Rost B. 1999. Twilight zone of protein sequence alignments. Protein Eng 12:85–94.

18. Tibshirani R. 1996. Regression shrinkage and selection via the lasso. J R Stat Soc Ser B Stat Methodol 58:267–288.

19. Winkler AM, Webster MA, Vidaurre D, Nichols TE, Smith SM. 2015. Multi-level block permutation. Neuroimage 123:253–268.

20. Salam N, Xian W-D, Asem MD, Xiao M, Li W-J. 2021. From ecophysiology to cultivation methodology: filling the knowledge gap between uncultured and cultured microbes. Mar Life Sci Technol 3:132–147.

21. Chung SY, Subbiah S. 1996. A structural explanation for the twilight zone of protein sequence homology. Structure 4:1123–1127.

22. Ghatak S, King ZA, Sastry A, Palsson BO. 2019. The y-ome defines the 35% of Escherichia coli genes that lack experimental evidence of function. Nucleic Acids Res 47:2446–2454.

23. Bernard G, Pathmanathan JS, Lannes R, Lopez P, Bapteste E. 2018. Microbial dark matter investigations: how microbial studies transform biological knowledge and empirically sketch a logic of scientific discovery. Genome Biol Evol 10:707–715.

24. Prakash O, Parmar M, Vaijanapurkar M, Rale V, Shouche YS. 2021. Recent trend, biases and limitations of cultivation-based diversity studies of microbes. FEMS Microbiol Lett 368:fnab118.

25. Almeida A, Nayfach S, Boland M, Strozzi F, Beracochea M, Shi ZJ, Pollard KS, Sakharova E, Parks DH, Hugenholtz P. 2021. A unified catalog of 204,938 reference genomes from the human gut microbiome. Nat Biotechnol 39:105–114.

26. Godzik A. 2011. Metagenomics and the protein universe. Curr Opin Struct Biol 21:398– 403.

27. Solden L, Lloyd K, Wrighton K. 2016. The bright side of microbial dark matter: lessons learned from the uncultivated majority. Curr Opin Microbiol 31:217–226.

28. Escudeiro P, Henry CS, Dias RP. 2022. Functional characterization of prokaryotic dark matter: the road so far and what lies ahead. Curr Res Microb Sci 3:100159.

29. Pavlopoulos GA, Baltoumas FA, Liu S, Selvitopi O, Camargo AP, Nayfach S, Azad A, Roux S, Call L, Ivanova NN. 2023. Unraveling the functional dark matter through global metagenomics. Nature 622:594–602.

30. Rodrigues DF, Ivanova N, He Z, Huebner M, Zhou J, Tiedje JM. 2008. Architecture of thermal adaptation in an Exiguobacterium sibiricum strain isolated from 3 million year old permafrost: a genome and transcriptome approach. BMC Genomics 9:1–17.

31. Sunagawa S, Acinas SG, Bork P, Bowler C, Eveillard D, Gorsky G, Guidi L, Iudicone D, Karsenti E. 2020. Tara Oceans: towards global ocean ecosystems biology. Nat Rev Microbiol 18:428–445.

32. Berney M, Cook GM. 2010. Unique flexibility in energy metabolism allows mycobacteria to combat starvation and hypoxia. PloS One 5:e8614.

33. Szenk M, Dill KA, de Graff AM. 2017. Why do fast-growing bacteria enter overflow metabolism? Testing the membrane real estate hypothesis. Cell Syst 5:95–104.

34. Burstein D, Sun CL, Brown CT, Sharon I, Anantharaman K, Probst AJ, Thomas BC, Banfield JF. 2016. Major bacterial lineages are essentially devoid of CRISPR-Cas viral defence systems. Nat Commun 7:10613.

35. Attrill EL, Łapińska U, Westra ER, Harding SV, Pagliara S. 2023. Slow growing bacteria survive bacteriophage in isolation. ISME Commun 3:95.

36. Boyd ES, Thomas KM, Dai Y, Boyd JM, Outten FW. 2014. Interplay between Oxygen and Fe–S Cluster Biogenesis: Insights from the Suf Pathway. Biochemistry 53:5834–5847.

37. Mand TD, Metcalf WW. 2019. Energy Conservation and Hydrogenase Function in Methanogenic Archaea, in Particular the Genus*Methanosarcina*. Microbiol Mol Biol Rev 83.

38. Mee MT, Collins JJ, Church GM, Wang HH. 2014. Syntrophic exchange in synthetic microbial communities. Proc Natl Acad Sci 111.

39. Norton JM, Klotz MG, Stein LY, Arp DJ, Bottomley PJ, Chain PSG, Hauser LJ, Land ML, Larimer FW, Shin MW, Starkenburg SR. 2008. Complete Genome Sequence of *Nitrosospira multiformis*, an Ammonia-Oxidizing Bacterium from the Soil Environment. Appl Environ Microbiol 74:3559–3572.

40. Stewart EJ. 2012. Growing Unculturable Bacteria. J Bacteriol 194:4151–4160.

41. Xavier JC, Patil KR, Rocha I. 2017. Integration of biomass formulations of genome-scale metabolic models with experimental data reveals universally essential cofactors in prokaryotes. Metab Eng 39:200–208.

42. Zheng M, Wen L, He C, Chen X, Si L, Li H, Liang Y, Zheng W, Guo F. 2024. Sequencing-guided re-estimation and promotion of cultivability for environmental bacteria. Nat Commun 15:9051.

43. Guillen MN, Rosener B, Sayin S, Mitchell A. 2021. Assembling stable syntrophic Escherichia coli communities by comprehensively identifying beneficiaries of secreted goods. Cell Syst 12:1064–1078.

44. Davis KER, Joseph SJ, Janssen PH. 2005. Effects of Growth Medium, Inoculum Size, and Incubation Time on Culturability and Isolation of Soil Bacteria. Appl Environ Microbiol 71:826–834.

45. Overmann J. 2006. Principles of enrichment, isolation, cultivation and preservation of prokaryotes. Prokaryotes 1:80–136.

46. Hessler T, Huddy RJ, Sachdeva R, Lei S, Harrison ST, Diamond S, Banfield JF. 2023. Vitamin interdependencies predicted by metagenomics-informed network analyses and validated in microbial community microcosms. Nat Commun 14:4768.

47. Alain K, Querellou J. 2009. Cultivating the uncultured: limits, advances and future challenges. Extremophiles 13:583–594.

48. Hamm JN, Erdmann S, Eloe-Fadrosh EA, Angeloni A, Zhong L, Brownlee C, Williams TJ, Barton K, Carswell S, Smith MA, Brazendale S, Hancock AM, Allen MA, Raftery MJ, Cavicchioli R. 2019. Unexpected host dependency of Antarctic Nanohaloarchaeota. Proc Natl Acad Sci 116:14661–14670.

49. Zengler K, Zaramela LS. 2018. The social network of microorganisms—how auxotrophies shape complex communities. Nat Rev Microbiol 16:383–390.

50. Chitty JL, Fraser JA. 2017. Purine acquisition and synthesis by human fungal pathogens. Microorganisms 5:33.

51. Moffatt BA, Ashihara H. 2002. Purine and pyrimidine nucleotide synthesis and metabolism. Arab BookAmerican Soc Plant Biol 1.

52. Schwengers O, Hain T, Chakraborty T, Goesmann A. 2019. ReferenceSeeker: rapid determination of appropriate reference genomes. BioRxiv 863621.

53. Ondov BD, Treangen TJ, Melsted P, Mallonee AB, Bergman NH, Koren S, Phillippy AM. 2016. Mash: fast genome and metagenome distance estimation using MinHash. Genome Biol 17:132.

54. Buchfink B, Reuter K, Drost H-G. 2021. Sensitive protein alignments at tree-of-life scale using DIAMOND. Nat Methods 18:366–368.

55. Babjac A, Royalty T, Steen AD, Emrich SJ. 2022. A comparison of dimensionality reduction methods for large biological data, p. 1–7. *In* Proceedings of the 13th ACM International Conference on Bioinformatics, Computational Biology and Health Informatics. ACM, Northbrook Illinois.

## Supplemental References

1. Chawla NV, Bowyer KW, Hall LO, Kegelmeyer WP. 2002. SMOTE: synthetic minority over-sampling technique. J Artif Intell Res 16:321–357.

2. Queen O, Emrich SJ. 2021. LASSO-based feature selection for improved microbial and microbiome classification, p. 2301–2308. *In* 2021 IEEE International Conference on Bioinformatics and Biomedicine (BIBM). IEEE.

3. Babjac A, Royalty T, Steen AD, Emrich SJ. 2022. A comparison of dimensionality reduction methods for large biological data, p. 1–7. *In* Proceedings of the 13th ACM International Conference on Bioinformatics, Computational Biology and Health Informatics. ACM, Northbrook Illinois.

4. Fonti V, Belitser E. 2017. Feature selection using lasso. VU Amst Res Pap Bus Anal 30:1– 25.

5. Fisher A, Rudin C, Dominici F. 2019. All models are wrong, but many are useful: Learning a variable’s importance by studying an entire class of prediction models simultaneously. J Mach Learn Res 20:1–81.

6. Pedregosa F, Varoquaux G, Gramfort A, Michel V, Thirion B, Grisel O, Blondel M, Prettenhofer P, Weiss R, Dubourg V. 2011. Scikit-learn: Machine learning in Python. J Mach Learn Res 12:2825–2830.

7. Huang J, Ling CX. 2005. Using AUC and accuracy in evaluating learning algorithms. IEEETrans Knowl Data Eng 17:299–310.

